# Host response to cholestyramine can be mediated by the gut microbiota

**DOI:** 10.1101/2020.12.08.416487

**Authors:** Nolan K. Newman, Philip M. Monnier, Richard R. Rodrigues, Manoj Gurung, Stephany Vasquez-Perez, Kaito A. Hioki, Renee L. Greer, Kevin Brown, Andrey Morgun, Natalia Shulzhenko

## Abstract

The gut microbiome has been implicated as a major factor contributing to metabolic diseases as well as being contributors to the response to drugs used for the treatment of such diseases. In this study, using a diet-induced obesity mouse model, we tested the effect of cholestyramine, a bile acid sequestrant, on the murine gut microbiome and mammalian metabolism. We also explored the hypothesis that some beneficial effects of this drug on systemic metabolism can be attributed to alterations in gut microbiota. First, we demonstrated that cholestyramine can decrease glucose and epidydimal fat levels. Next, while investigating gut microbiota we found increased alpha diversity of the gut microbiome of cholestyramine-treated mice, with fourteen taxa showing restoration of abundance to levels resembling those in mice fed with a control diet. Analyzing expression of genes known to be regulated by cholestyramine (including Cyp7a1), we confirmed the expected effect of this drug in the liver and ileum. Finally, using a transkingdom network analysis we inferred *Acetatifactor muris* and *Muribaculum intestinale* as potential mediators/modifiers of cholestyramine effects on the mammalian host. In addition, *A. muris* correlated positively with glucagon (Gcg) expression in the ileum and negatively correlated with small heterodimer partner (Shp) expression in the liver. Interestingly, *A. muris* also correlated negatively with glucose levels, further indicating the potential probiotic role for *A. muris*. In conclusion, our results indicate the gut microbiome has a role in the beneficial effects of cholestyramine and suggest specific microbes as targets of future investigations.

## Introduction

As of the year 2016, hypercholesterolemia (excessive amounts of cholesterol in the blood) is prevalent in more than 29.4% of American adults over the age of 20 (National Center for Health Statistics, 2018), up from 25% in the year 2002. Individuals with familial hypercholesterolemia are at a higher risk of developing cardiovascular diseases (Masana et al., 2019) and 60-70% of patients with Type 2 Diabetes Mellitus (T2D) also have dyslipidemia (Parhofer, 2015). An increase in foods rich in animal fats and sugars, termed a “Western diet” (WD), has led to an increasing number of people with hypercholesterolemia and T2D, making it increasingly important to understand how anti-hypercholesterolemic drugs act in the body. Recently, it has become clear that both antidiabetics (Whang et al., 2019; Gu et al., 2017; Gurung et al., 2020) and anti-hypercholesterolemic drugs cause alterations in the host microbiome. Atorvastatin, a commonly used HMG-CoA reductase inhibitor, has been shown to alter the gut microbiome of rats fed a high-fat diet (Khan et al., 2018). Other anti-hypercholesterolemic drugs, such as cholestyramine, act by binding to bile acids in the gut and preventing their absorption, increasing excretion of bile acids in the stool (Scaldaferri et al., 2013). This, in turn, increases cholesterol metabolism in the liver, thus reducing blood cholesterol levels. While cholestyramine has been shown to improve serum glucose levels in rats (Chen et al., 2010), a common marker of diabetes, its interaction with the gut microbiome has not been well established. A study published in 1975 (Williams et al., 1975) showed a decrease in anaerobic bacteria after one week of cholestyramine but comprehensive analysis of microbiome was not possible back then. More recently, it was reported that treatment with cholestyramine results in an increased abundance of a pathogen *Clostridium difficile* in the intestine (Buffie et al., 2014). Due to the interrelationships between bile acids and the gut microbiome (Ridlon et al., 2014), it has been hypothesized that cholestyramine’s bile acid sequestration mechanism would cause changes in the gut microbiome composition (Jia et al., 2018). We sought to determine whether cholestyramine would alter the composition of the gut microbiome in mice fed a high animal fat, high sugar diet. Our results demonstrate that in WD-fed animals, cholestyramine treatment results in large shifts in the gut microbiome composition, especially in the Lachnospiraceae and Ruminococcaceae families, both of which are from the order Clostridiales. Furthermore, through reconstruction and interrogation of a transkingdom network, we identified two microbial species, *Acetatifactor muris* (family Lachnospiraceae, order Clostridiales) and *Muribaculum intestinale* (family Porphyromonadaceae, order Bacteroidales) that likely play a part in facilitating the response to cholestyramine. Altogether, these results implicate that cholestyramine at least partially acts through altering the gut microbiome and its mechanism may be dependent on abundances of the species *A. muris* and *M. intestinale*.

## Results

### Cholestyramine restores systemic parameters associated with metabolic disease

To mimic the disease treatment, mice were first fed a WD for 8 weeks to induce T2D followed by 8 weeks of a WD with added cholestyramine (WD+Ch) and systemic parameters associated with T2D were measured (**Supplementary figure 1**). The control groups were either fed a Western diet (WD) or normal diet (ND) for 18 weeks. Mice on a WD demonstrated an increase in 11 parameters associated with metabolic disease seen in diabetic individuals, when compared to the ND group (**Figure 1a,c, Supplementary table 1**). Eight of these parameters decreased, six of which were significant (one-sided Mann-Whitney U FDR < 0.1), in the WD+Ch group. In particular, parameters associated with glucose metabolism (i.e. fed glucose levels, fasting glucose levels, and glucose levels at various time points after injecting glucose; **Figure 1a,b, Supplementary figure 2**), were decreased by cholestyramine (WD+Ch), along with epididymal fat mass. Although not statistically significant, serum cholesterol levels dropped as expected with cholestyramine treatment (p-value = 0.11).

**Figure 1.**
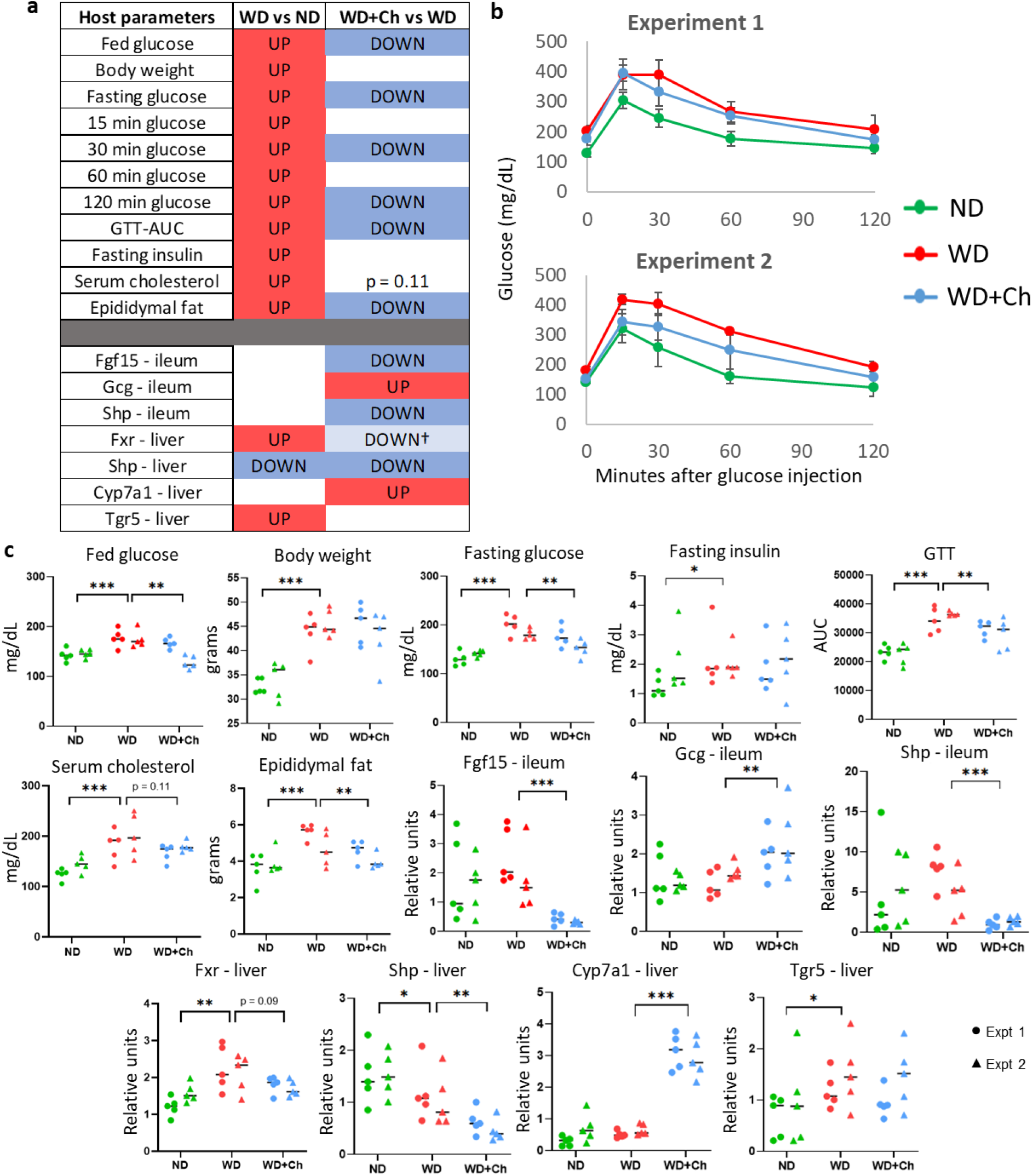
a) Summary table of significantly changed host parameters (phenotypes: one-tailed Mann-Whitney U FDR < 0.1); genes: two-tailed Mann-Whitney U FDR < 0.1) and their fold change directions (red = up, blue = down). b) GTT-AUC plots demonstrating the changes in blood glucose levels over the first 2 hours after glucose injection. c) Dot plots of phenotypes and genes that demonstrated significant differences in fold changes. Circles represent experiment 1 and triangles represent experiment 2. *,**,*** = p-value < 0.05, 0.01, or 0.001 respectively. † = p-value < 0.1.

### Cholestyramine changes the expression of genes involved in bile acid metabolism

Due to cholestyramine’s mechanism of action (sequestration of bile acids and limiting reabsorption), we hypothesized that treatment with cholestyramine would cause a change in expression of genes involved in the bile acid metabolism pathway (Chiang & Ferrell, 2018; Kliewer & Mangelsdorf, 2016). The group of mice on a WD exhibited increased expression of Fxr (Nr1h4) and Tgr5 and decreased expression of Shp in the liver (**Figure 1a, 1c**, two-sided Mann-Whitney U FDR < 0.1). Of the genes measured in the bile acid metabolism pathway, three in the ileum were altered by treatment (Fgf15, Gcg, and Shp). Fgf15 and Shp in the ileum decreased expression with treatment while Gcg in the ileum increased (**Figure 1a,c**). Only Shp and Cyp7a1 were significantly altered in the liver after treatment, which exhibited decreased and increased expression, respectively. Gene expression of Fxr in the liver showed a trend toward decreased expression (p-value = 0.09).

### Cholestyramine alters the gut microbiome composition

Previous research indicated that the gut microbiome can influence ileal gene expression levels (Larsson et al., 2012) and the host’s ability to metabolize glucose (Rodrigues et al., 2017). This led us to ask whether changes in the gut microbiome could be in part responsible for the restoration of metabolic parameters observed following treatment with cholestyramine. We thus performed analysis of the microbial 16S rRNA genes in the ileum samples of the same mice used in **Figure 1**.

Amplicon Sequence Variants (ASVs) in the cholestyramine-treated samples separated from those in the ND and WD samples primarily on PC1, which accounted for 48.7% and 63.3% of the variability seen in experiments 1 and 2, respectively (**Figure 2a**). All groups demonstrated a significant difference in microbial composition in both experiment 1 (ANOSIM R-value 0.957, *p*-value < 0.001) and experiment 2 (ANOSIM R-value 0.966, *p*-value < 0.001). The ND and WD samples separated mainly on PC2, which accounted for 24% and 16.9% of the differences seen in experiments 1 and 2, respectively. All groups clustered distinctly and there was no overlap between groups. Comparisons between WD and ND groups, as well as WD+Ch and WD groups can be seen in **Supplementary figure 3**. Alpha diversity of families increased significantly in the WD+Ch group compared to the WD group (p-value < 0.001, **Figure 2b**). Looking deeper into the differences between WD and WD+Ch groups, we found that, across both experiments, treatment with cholestyramine decreased the abundance of microbiota in four families (Ruminococcaceae, S24-7, Coriobacteriaceae, and Dehalobacteriaceae) while increasing the abundance of four other families (Anaeroplasmataceae, Lachnospiraceae, Turicibacteraceae, and Clostridiaceae, **Figure 2c, Supplementary figure 5, Supplementary table 2**).

**Figure 2.**
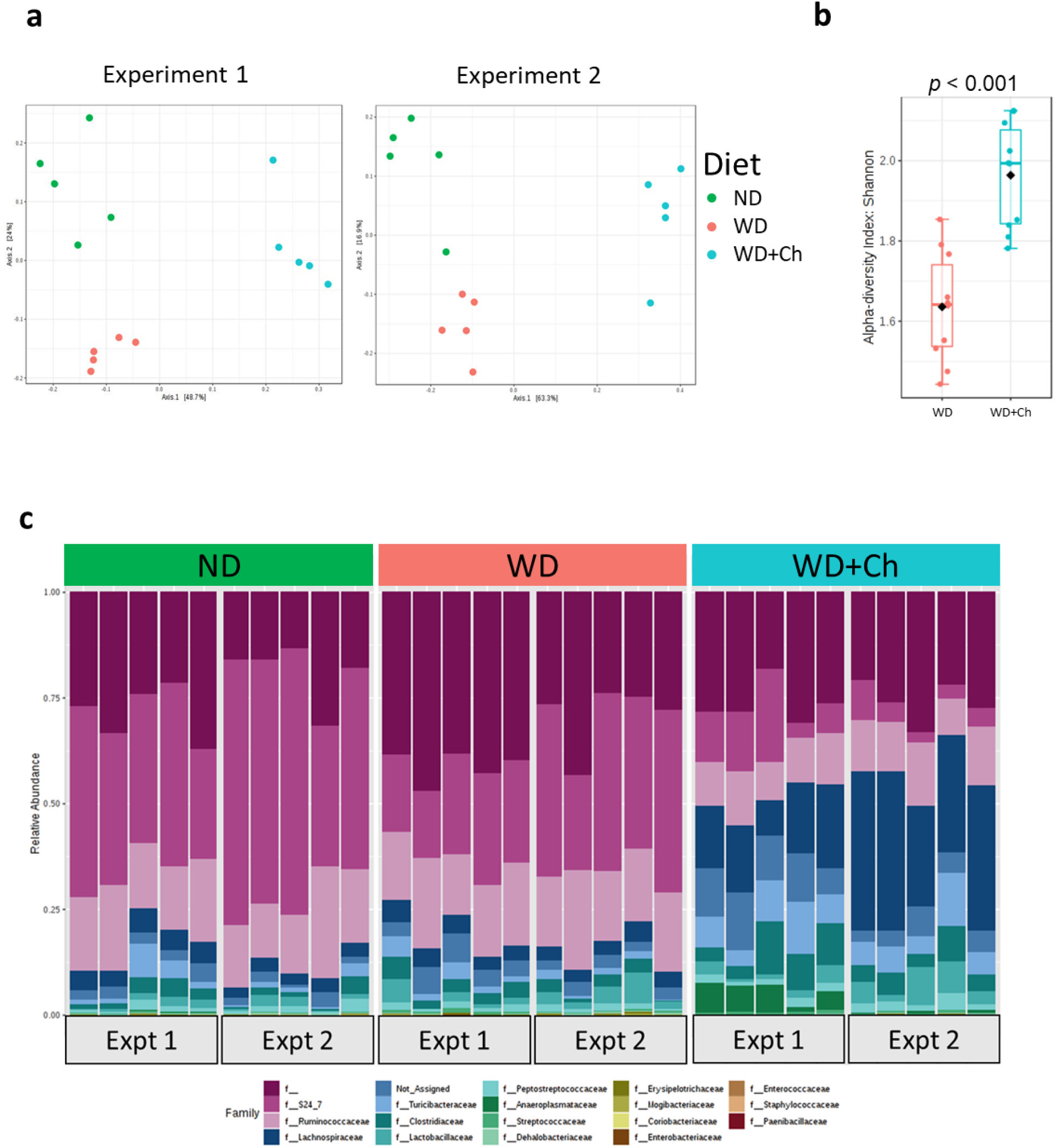
a) PCoA plots demonstrating there is a shift in microbial composition in WD fed mice administered cholestyramine (blue) from that of the WD fed, cholestyramine-free group (red) and mice administered a normal chow diet (green). ANOSIM R-values for Experiment 1 and Experiment 2 are 0.957 and 0.966, respectively, and each has a p-value < 0.001. b) Family level alpha diversity is significantly higher in the WD+Chol group (Mann-Whitney *p-value* < 0.001) c) Relative abundances of the families found in each individual.

To estimate which of the ASVs contributed to the observed restoration of the 6 metabolic parameters, we classified ASVs into three distinct groups (**Figure 3**). Group 1 refers to those ASVs that demonstrated opposite fold change direction between WD and WD+Ch treatments, meaning those that were increased (or decreased) by WD and decreased (or increased) in the WD+Ch group. Thus, the abundances of these ASVs were reversed towards normal levels by treatment with cholestyramine. Among the 14 ASVs found, 13 belonged to the order Clostridiales while one belonged to Bacteroidales. Group 2 contained those ASVs that were changed in only the WD+Ch treatment group, indicating these changes were independent of diet. Of the 31 ASVs, the majority were also Clostridiales. Finally, group 3 contained 17 ASVs that changed in the same direction in both the WD and WD+Ch compared to the ND group. The fact that these were not dependent on cholestyramine indicates they are potentially non-causal microbes for the response to cholestyramine.

Although we identified 14 ASVs that are restored by cholestyramine treatment, those that regulate the host parameters cannot be distinguished by changes in their abundances alone. Thus, to identify potential causal microbes, we turned to reconstructing a co-variation network between the phenotypes, genes, and ASVs in the WD+Ch treated group.

**Figure 3.**
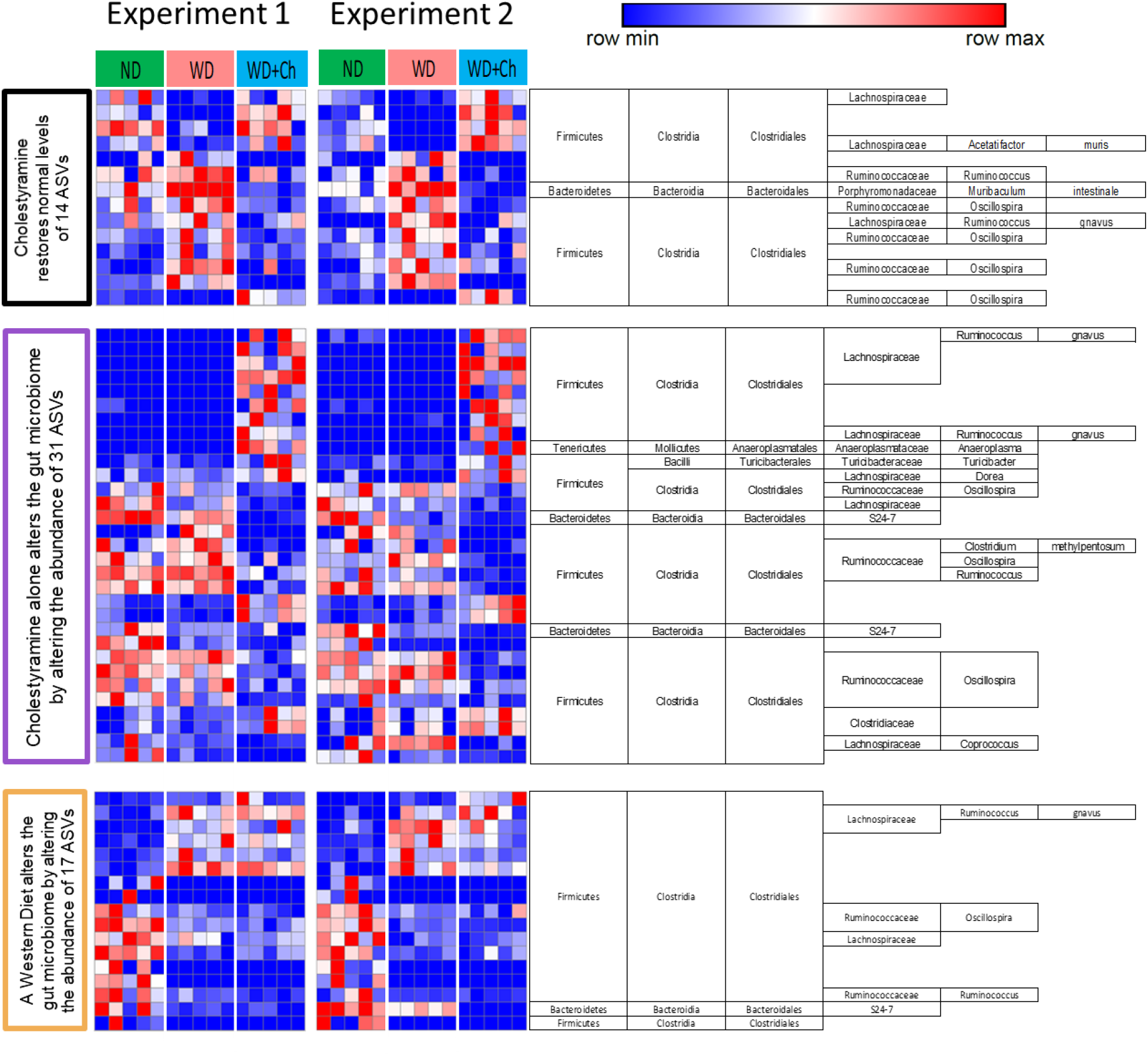
Group 1: 14 ASV abundances are restored after the introduction of cholestyramine to a WD. Group 2: The abundance of 31 ASVs are altered by cholestyramine alone. Group 3: 17 ASVs are altered by a WD, regardless of treatment with cholestyramine. The color of the labels for each heatmap corresponds to the outlines of nodes in the transkingdom network (Figure 4). Blue and red colors indicate decreased and increased ASV abundances, respectively. Rows (ASVs) of heatmaps were clustered first by abundance in experiment 1 and the same order was used for experiment 2. Significant ASVs were required to be consistent in fold-change direction across both experiments, Mann-Whitney *p*-value < 0.05).

### Transkingdom network reconstruction identifies *Acetatifactor muris* and *Muribaculum intestinale* as potential contributors to response to cholestyramine

Networks are commonly used in systems biology to identify relationships between hosts and the gut microbiome (Morgun et al., 2015). The network we constructed herein included interactions (correlations) between the host parameters (both metabolic parameters and genes) and the gut microbiome of the cholestyramine treated mice (**Figure 4a**). 84 nodes and 154 edges were retained in our network after filtering out nodes and edges that did not satisfy statistical parameters (see materials and methods) and causality principles (correlation inequalities, Yambartsev et al., 2016; VanderWeele & Robins, 2010). The network was composed of 73 ASVs (53 of which were in the main component), 6 metabolic parameters, and 5 genes. The main component of the network included major parameters of glucose metabolism and 3 genes, 2 of which were from ileum. As expected for a biological network, the relationship between the number of nodes and number of edges followed a power-law distribution (**Supplementary figure 6**).

**Figure 4.**
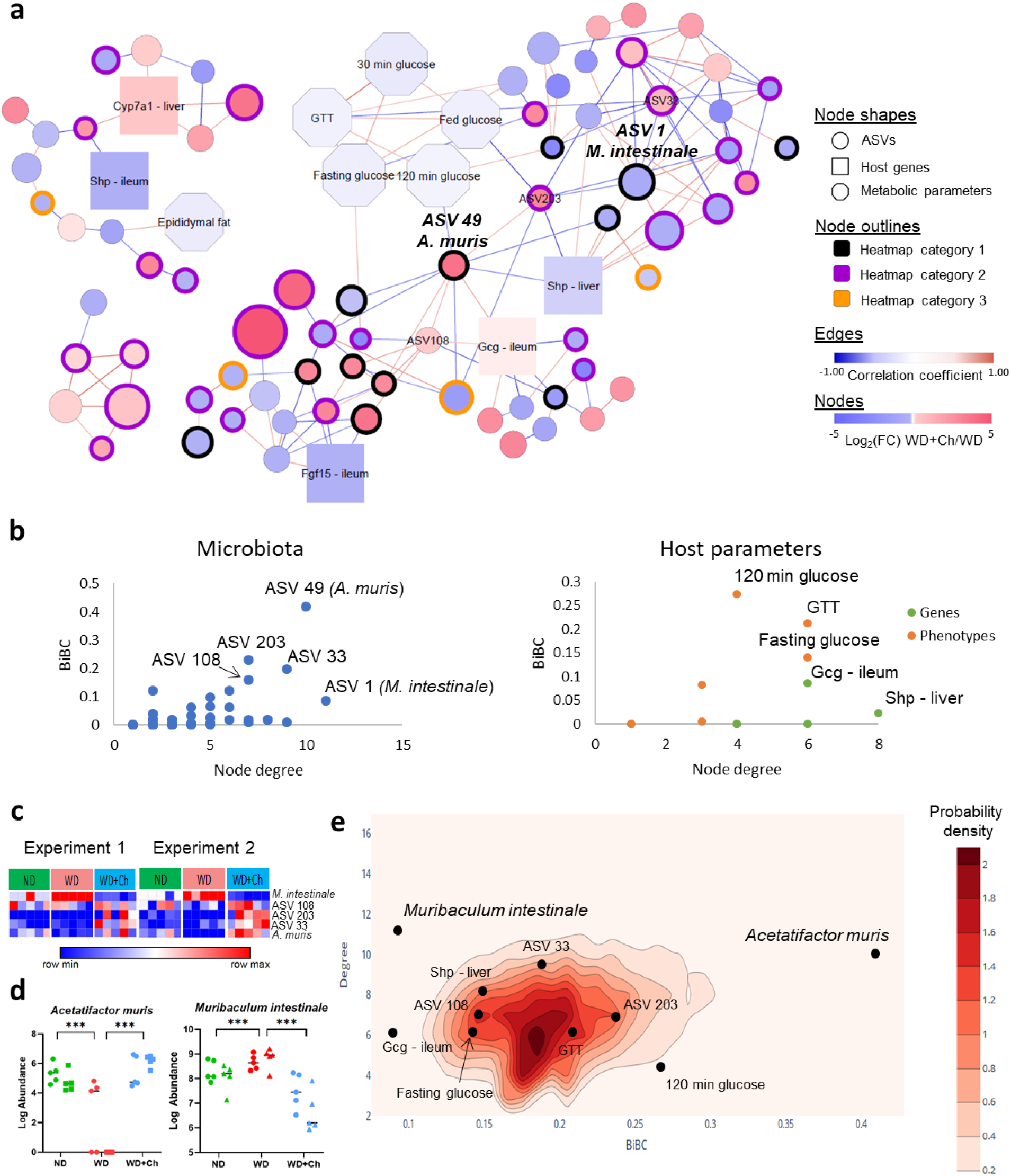
a) Transkingdom network showing correlations between genes, phenotypes, and ASVs. ASV outlines reflect the heatmap group that microbe belongs to (heatmap #1 = black, #2 = purple, #3 = orange). Edges represent significant Spearman correlations (Spearman *p* < 0.05) and are colored based on Spearman’s rho coefficients (blue = negative correlation, red = positive correlation). Nodes are colored based on fold-change direction (red = up, blue = down) of the node between the Cholestyramine+WD group and the WD group (Mann-Whitney U p-value < 0.05). ASV size is based on log-transformed abundance b) BiBC-degree distribution of all nodes, the highest of which are marked. c) Heatmap of abundances of top five ASVs ranked on BiBC-degree distribution d) Abundances of the top two ASVs (*A. muris* and *P. catoniae*). e) 2D-contour histogram of the nodes with highest BiBC and degree from 10,000 randomly generated networks. Darker areas indicate higher probabilities of randomly finding a node with that degree and BiBC.

To determine which microbiota members were most likely to have a regulatory effect on the host response to cholestyramine, we focused on two key node properties: degree (the number of nodes a given node connects to) and bipartite betweenness centrality (BiBC). BiBC is similar to betweenness centrality but BiBC splits the network up into groups (i.e. microbiota and metabolic parameters; Dong et al., 2015). BiBC is essentially a measure of the “bottleneck-ness” of a node. The BiBC of a node counts how many times that node lies on the shortest paths between all possible pairs of nodes formed by selecting one node from the first group and one node from the second group. A microbe with a high BiBC and large degree is one that likely plays a role in host response to cholestyramine. By comparing BiBC and degree of each parameter (using microbiota and host metabolic parameters as our two groups for the BiBC analysis), we found two potentially important ASVs (**Figure 4b-d**). ASV 49, which had both a large BiBC and degree, positively correlated with Gcg expression in the ileum and negatively correlated with both Shp in the liver and glucose 120 minutes in the glucose tolerance test. ASV 1, which did not have a large BiBC but did have a high degree, primarily correlated with other ASVs.

After using BLAST (Boratyn et al., 2013) to identify these ASVs at the species level, the closest match to ASV 49 was *Acetatifactor muris* (91.9% identical) and the closest match to ASV 1 was *Muribaculum intestinale* (93.3% identical). This indicates *A. muris*, a member of the family Lachnospiraceae, to be a likely regulator of the host metabolic response to cholestyramine and *M. intestinale*, a member of the family Porphyromonadaceae, to be a microbe that communicates with many other microbes in the gut. *A. muris* was found to belong to the first heatmap group (**Figure 3**) and had its abundance increased in the WD+Ch group from that of the WD group. *M. intestinale* also belonged to the first group but had its abundance decreased in response to cholestyramine treatment. Using the same methods, we discovered Gcg in the ileum and Shp in the liver were important genes in regulating the host-microbiome interactions.

To further validate our findings that *A. muris* and *M. intestinale* were important players in the host response to cholestyramine, we generated an ensemble of 10,000 random networks (see materials and methods) and compared the BiBC and degree of the two microbes in the real network to the nodes with the largest BiBC and degree in the randomly generated networks (**Figure 4e**). As predicted, the actual BiBC and degrees of *A. muris* and *M. intestinale* were atypical of those found in a random network (nodes with degrees and BiBC values that were high were extremely rare in the random ensemble). Meanwhile, those nodes that had a smaller BiBC and degree were clustered more towards the center of the BiBC-degree distribution, indicating they were not as important in the functional role of cholestyramine on the host.

## Discussion

From our study, it is clear that cholestyramine has more than just anti-hypercholesterolemic effects on the host. Consistent with findings from other studies, it can also help restore the ability of obese mice to metabolize glucose (Chen et al., 2010; Garg & Grundy, 1994). Furthermore, other bile acid sequestrants such as colesevelam, colestilan, and sevelamer have been repeatedly shown to improve glucose levels in diseased individuals (Bays et al., 2008, Fonseca et al., 2008; Kondo et al., 2010; Brønden et al., 2018). While it has previously been shown cholestyramine can exert its anti-hyperglycemic effects via increasing ileal expression of Gcg (which encodes for preproglucagon, a precursor to GLP-1; White & Grady, 1986), the exact mechanism for this increase remains unknown. Our discovery that *A. muris* positively correlates with Gcg expression in the ileum indicates that cholestyramine may act on Gcg expression via an increase in *A. muris*.

*A. muris* was first identified in the caecum of obese mice and was noted to be a producer of both acetate and butyrate (Pfeiffer et al., 2012). Previous research has demonstrated the importance of acetate in reducing fat accumulation in the liver and improving host response to insulin (Yamashita et al., 2007). Similarly, butyrate administration in the diet corresponds with increased insulin sensitivity in mice (Gao et al., 2009) and sodium butyrate can reduce cholesterol, LDL, VLDL, and plasma triglyceride in diabetic rats (Khan & Jena, 2016). Others have proposed that a main role of butyrate is to reduce expression of genes involved in the synthesis of intestinal cholesterol (Alvaro et al., 2008), which could in part explain the increased abundance of *A. muris* and decreased levels of cholesterol in the WD+Ch.

The results of our transkingdom network indicate that *M. intestinale* may also play a role in crosstalk between microbiota in response to cholestyramine, but because of its relatively low BiBC value, *M. intestinale* does not appear to have as direct of a regulatory role as *A. muris*.

A previous study has shown that *M. intestinale* decreased in response to a high-fat diet (Do et al., 2018). In contrast, *M. intestinale* increased in our WD group and decreased after treatment with cholestyramine. This discrepancy may be due to differences in diet composition. In addition, the 16S sequences of ASV 1 and ASV 49 were not 100% identical to their predicted species, so the real identity may be a closely related species.

The changes of genes associated with bile acid metabolism in the cholestyramine group (**Figure 1c**) is what one would expect to see with a bile acid sequestrant (**Figure 5**). Specifically, the decreased availability of bile acids in the ileum (through sequestration by cholestyramine) decreased activity of ileal farnesoid X receptor (Fxr, Nr1h4), which in turn would lead to decreased expression of ileal small heterodimer partner (Shp, a.k.a. Nr0b2; Neimark et al., 2004) and fibroblast growth factor 15 (Fgf15, the ortholog of FGF19 in humans), as Fxr is a positive regulator of Fgf15. Lower expression of Fgf15 in the ileum would lead to lower expression of Shp in the liver (Kliewer & Mangelsdorf, 2016; Keeley & Walters, 2016). Since hepatic Shp normally inhibits Cyp7a1, Cyp7a1 expression increases following the inhibition of Shp with cholestyramine, (Goodwin et al., 2016; Zhang & Klaasen, 2010). Hence, a decreased expression of ileal Fgf15 via cholestyramine administration leads to an increased expression of Cyp7a1.

**Figure 5.**
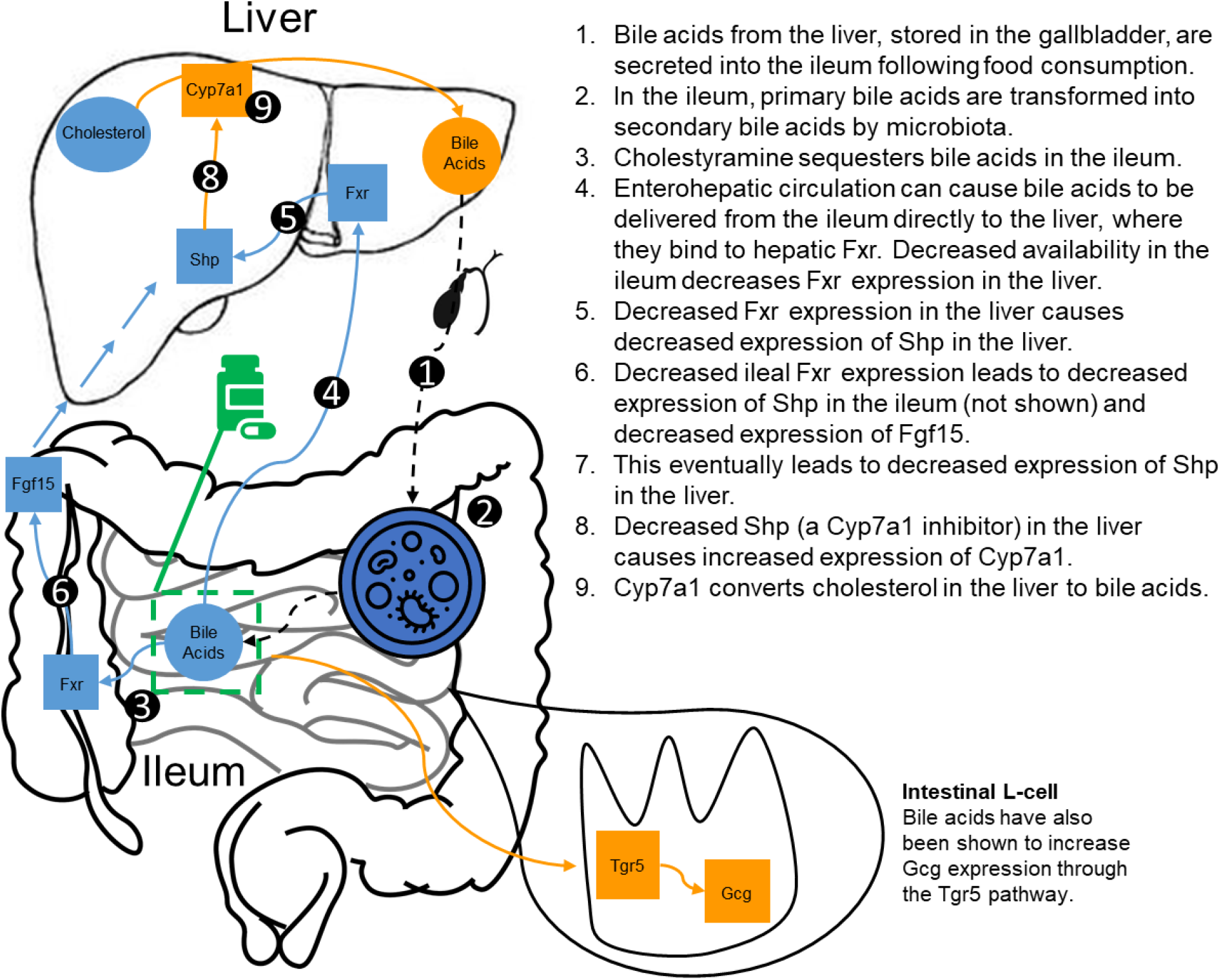
Representation of the expected fold change direction of genes involved in the bile acid metabolism pathway. Figure represents what is expected to happen following cholestyramine treatment (orange = increase, blue = decrease)

Another non-mutually exclusive pathway for bile acid metabolism through Cyp7a1 mediation has been proposed, in which enterohepatic circulation causes bile acids from the ileum to be transported to the liver, where they bind to activate Fxr, which then inhibits Shp (Chiang, 2017).

In the cholestyramine group, the ileal Fxr expression tended to be lower. The downstream target of Fxr, Fgf15, exhibited decreased expression in the ileum, resulting in decreased Shp expression (both ileal and liver), and thus a higher expression of Cyp7a1. Interestingly, while we did not observe an increase in ileal Tgr5, its downstream target (Wan et al., 2018), Gcg was increased. This seems to indicate that another mechanism is responsible for the increase in Gcg expression in cholestyramine-treated mice. Such mechanism might involve changes in microbiota, in particular, *A. muris*, which is predicted by our analysis as a positive regulator of Gcg gene expression. This prediction could be tested in future studies. It has also been hypothesized that bile acid sequestrants increase fatty acids in the ileum through decreased micelle production (Sonne et al., 2014). In turn, the increased availability of fatty acids may cause increases in Gcg expression (Hirasawa et al., 2004). Although we did not see significantly lower levels of Fxr expression in the ileum, it has been previously shown that colesevelam, another bile acid sequestrant, can inhibit L-cell Fxr expression, which in turn can act to increase Gcg expression (Trabelsi et al., 2017).

To the best of our knowledge, our study is the first to report large changes in the microbiome following cholestyramine treatment. Our results are consistent with other studies showing that statins and bile acid sequestrants can modify the microbial composition of the gut (Caparrós-Martín et al., 2017; Khan et al., 2018; Fuchs et al., 2018). Considering the crucial role of the gut microbiome in metabolizing bile acids (including deconjugation and hydrolysis; Kriaa et al., 2019), the fact that fluctuations in bile acid sequestration levels lead to changes in the gut microbiome is not entirely surprising. It is well-known that bile acids are toxic to microbial cells, which requires the microbiota to chemically modify the bile acids as a defense mechanism (Ruiz et al., 2013). By binding to bile acids, cholestyramine and other bile acid sequestrants may decrease selective pressure on host microbiota that cannot deconjugate bile acids, resulting in the increase in alpha diversity we observed in the WD+Ch group.

## Conclusion

We have demonstrated that treatment of obese diabetic mice with the bile acid sequestrant cholestyramine lowers glucose levels possibly through interactions with the host microbiome. Mice treated with cholestyramine demonstrated significant changes in microbial composition as well as changes in the expression of genes involved in the bile acid metabolism pathway. Through reconstruction and interrogation of a data-driven transkingdom network, we were able to identify two potential players responsible for host response to cholestyramine, *Acetatifactor muris* and *Muribaculum intestinale*. The next step could be to perform experimental validation of these findings via the colonization of germ-free mice with each of the identified species to determine whether their presence is truly indicative of an individual’s response to cholestyramine. Our work has provided a foundation of knowledge for host-microbiota interactions under cholestyramine and can help in understanding the complex relationship between glucose, bile acid metabolism, and the gut microbiome.

## Materials and Methods

### Mice and Cholestyramine Treatment

7 weeks old, Specific Pathogen Free (SPF), C57BL/6 male mice were purchased from Jackson Laboratory (Bar Harbor, Maine) and housed in the controlled environment (12h daylight cycle) of the Laboratory Animal Resources Center (LARC) at Oregon State University with ad libitum access to food and water. After one week of acclimatization, mice were switched to either a Western diet (WD) D12451 containing 45% lard and 20% sucrose or a matching normal diet (ND) D12450K produced by Research Diets (New Brunswick, NJ). After 8 weeks, one group of mice was switched from WD alone to WD supplemented with 2% cholestyramine (20g/kg of diet, Sigma) for 10 weeks (**Supplementary Figure 1**). Two independent experiments (n = 5 per group per experiment) were performed. In experiment 2, mice were supplemented with cholestyramine for 11 weeks instead of 10. Body weight and feed intake were measured each week. An intraperitoneal glucose tolerance test was performed before the initiation of cholestyramine treatment and after the end of the experiment. Stool samples, whole fasting blood (with EDTA) and fasting serum (with DPP4 inhibitor) were collected via submandibular bleed. Following cholestyramine treatment, mice were sacrificed and tissues were collected.

### Glucose Tolerance Tests

Mice were fasted for 6 hours during the light phase with free access to water. A concentration of 2 mg/kg body weight glucose (Sigma-Aldrich) was injected intraperitoneally. Blood glucose was measured at 0 (immediately before glucose injection), 15, 30, 60 and 120 minutes with a Freestyle Lite glucometer (Abbot Diabetes Care).

### Serum Fasting Insulin

Mice were fasted for 6 hours with free access to water. Fasting blood was collected via submandibular bleed. Insulin in serum was measured with Mouse Insulin ELISA Kit (Crystal Chem) according to manufacturer’s protocol.

### Serum Cholesterol

Serum cholesterol was measured using a Cholesterol Assay Kit (Thermofisher) as per the manufacturer’s instructions. Non-fasted serum was used.

### Bacterial DNA Extraction and 16S rRNA Gene Library Preparation

For microbial measurements, fresh stool pellets were collected and immediately stored at −80°C. To extract microbial DNA, frozen fecal pellets were resuspended in 1.4 mL ASL buffer (Qiagen) and homogenized with 2.8 mm ceramic beads followed by 0.5 mm glass beads using an OMNI Bead Ruptor (OMNI International). DNA was extracted from the entire resulting suspension using a QiaAmp mini stool kit (Qiagen) according to the manufacturer’s protocol. DNA was quantified using the Qubit broad range DNA assay (LifeTechnologies). The V4 region of the 16s rRNA gene was amplified using universal primers (515f and 806r) as previously described (Greer et al., 2016). Individual samples were barcoded, pooled to construct the sequencing library, and then sequenced using an Illumina Miseq (Illumina, San Diego, CA) to generate paired-end 250 bp reads.

### Tissue collection, RNA Preparation and Gene Expression Analysis

Mouse ileum samples were homogenized in RLT buffer using the OMNItip tissue disruptor. RNA was then extracted using the QiaCube RNeasy kit with Qiashredder as per the manufacturer’s instructions. Mouse liver samples were homogenized in Trizol using the OMNItip tissue disruptor, and 200 ul of chloroform was added to each sample. Each tube was then centrifuged at 4°C and 12,000 g for 15 minutes. RNA was then extracted from the upper aqueous layer using the QiaCube RNeasy kit with Qiashredder, following the same protocol as for the ileum samples. On-column DNA digestion was used for both types of tissues. Concentrations of RNA were measured using the Quant-IT RNA BR Assay Kit (Life Technologies). Reverse transcription of RNA into cDNA was performed using the qScript cDNA Synthesis Kit (Quanta Biosciences) following the manufacturer’s protocol. Primer master mixes were prepared for Quantitative Real-Time PCR with the PerfeCTa SYBR Green PCR Kit (QuantaBio) with Polr2c as a housekeeping gene. Primers used for each gene are included in the below table. qRT-PCR was performed on the StepOnePlus Real-Time PCR system (Applied Biosystems) with the following protocol: samples were heated to 95°C and held for 30 seconds, followed by 40 cycles of 95°C for 3 seconds and 60°C for 20 seconds.

Primers for each gene, including the Polr2c housekeeping gene, are listed in the below table.

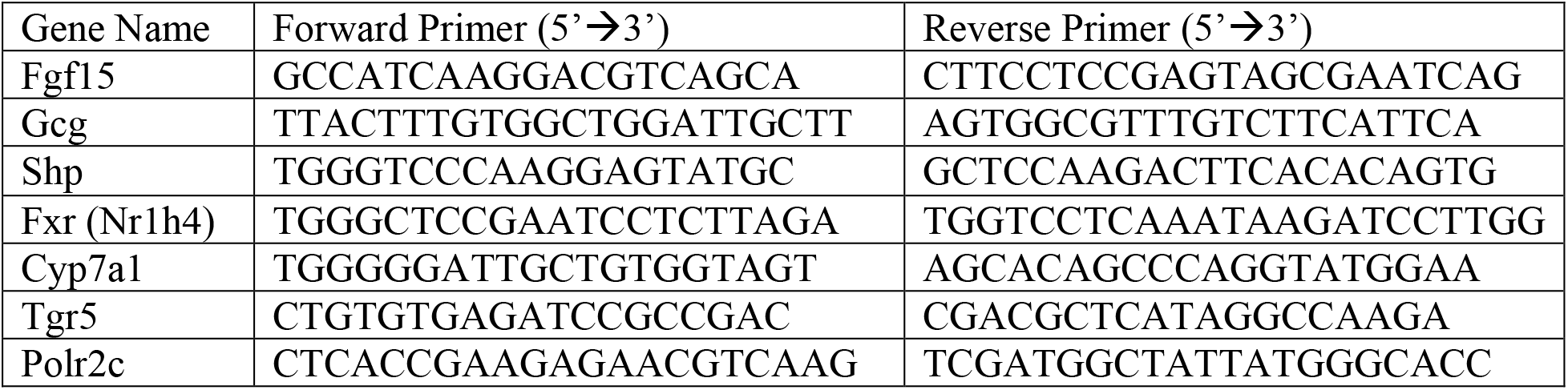

DNA concentrations of each gene relative to the housekeeping gene were calculated using the standard curve method. The gene expression standard for ileum and liver samples were created by pooling cDNA from all 30 mice samples. A 1:10 dilution of the standard was created to plot the standard curve. For each sample, 2^−Ct^ values were used to calculate DNA concentrations for each gene. DNA concentrations for each gene were calculated relative to the housekeeping gene for each sample.

### Statistical Analysis of Host Parameters

Phenotypes and genes (host parameters) were compared to their respective control groups. Host parameters were determined to be significant if: 1) fold change direction (+/−) was consistent across both experiments and 2) after median normalizing each experiment and pooling host parameters, the FDR corrected Mann-Whitney U p-value (one-tailed for metabolic parameters, two-tailed for genes) was less than 0.1.

### 16S rRNA Gene Sequencing Analysis

The QIIME 2 (Bolyen et al. 2019) bioinformatics pipeline (v. 2018.8.0) was used to demultiplex and quality filter the forward-end fastq files. Denoising was performed using DADA2 (Callahan et al., 2016). Taxonomy was assigned to the amplicon sequence variants (ASVs) using the naïve Bayes taxonomy classifier against the Greengenes 13_8 99% OTUs reference sequences (McDonald et al. 2012). Microbes with the frequency above 99.95% of cumulative abundance across all samples in an experiment were included in further analysis. Common ASVs within this threshold in both experiments were relativized per million and then quantile normalized.

For beta diversity analysis, the quantile normalized ASV tables were uploaded to MicrobiomeAnalyst (Chong et a., 2020; Dhariwal et al., 2017). Data was analyzed on MicrobiomeAnalyst per experiment. PCoA plots were generated using the Bray-Curtis dissimilarity distance method at the feature level and the analysis of similarities (ANOSIM) was calculated between treatment groups. Alpha diversity was measured using the Shannon diversity index and a Mann-Whitney U test was performed to compare the WD+Ch and WD groups. Stacked percent abundance plots were created at both the Family and Order level (**Supplementary figure 4**). Families that were significantly changed between the WD+Ch and WD groups were determined using a two-tailed Mann-Whitney U test (FDR < 0.05).

Prior to creating the heatmaps, raw ASVs were relativized per sample, then mean normalized across all samples (including both timepoints, but only the samples at the endpoint were visualized in the heatmaps). Following this, the data was filtered to contain only microbes that belonged to one of three categories: 1) The ASV significantly (two-tailed Mann-Whitney U test p-value < 0.1) increased (or decreased) in the WD group compared to the ND group, but significantly changed in the opposite direction in the WD+Ch groups compared to the WD group; 2) The ASV significantly increased (or decreased) in the WD+Ch group compared to ND and WD, but there was no difference when comparing the ND and WD group to one another; 3) The ASV significantly increased (or decreased) in the WD and WD+Ch group when compared to the ND group but there was no difference when comparing the WD and WD+Ch group to one another. The ASV was required to behave similarly across both experiments for it to be in the heatmap. After creating these three groups, heatmaps for experiment 1 were generated using the online Morpheus heatmap generator (https://software.broadinstitute.org/morpheus). Hierarchical clustering using the Euclidean distance metric and complete linkage was performed on rows and each row was scaled based on the maximum and minimum value of the row. Heatmaps for experiment 2 reflect the same row order as in experiment 1, meaning clustering was not performed on experiment 2.

### Network Reconstruction and Identification of Key Nodes

Differentially expressed genes and significant metabolic parameters were found by first checking for the same fold change direction (+/−) between both experiments, then median normalizing each parameter per experiment and pooling. Host parameters were considered to be significant if the FDR corrected one-tailed Mann-Whitney U p-value was less than 0.05 between the WD+Ch and WD groups. Significant ASVs were found similarly, although without median normalizing of the quantile normalized data prior to pooling. In the WD+Ch group, Spearman correlations were then performed between each metabolic parameter, gene, and ASV. Correlations were filtered based on the following criteria: 1) same direction of correlation (+/−) across both experiments prior to pooling, 2) Spearman correlation p-value < 0.1, 3) does not exhibit correlation inequalities (Yambartsev et al., 2016; Vanderweele and Robins, 2010). Networks were visualized using Cytoscape v3.7.2 (Shannon et al., 2003). The Python module NetworkX v2.2 (Hagberg et al., 2008) was used to calculate BiBC and degree between groups, as well as to randomly generate the 10,000 networks used in validating the BiBC and degree results. BiBC was calculated as previously described (Dong et al., 2015). Binomial random (Erdos-Renyi) networks were generated from the G(m,n) ensemble with m = 154 (to match the number of edges in the real network) and n = 84 (the number of nodes). No self-or multi-edges were allowed. The online tool Plotly (https://plot.ly/) was used to plot the 2D contour histogram of the BiBC-degree distribution. Probability density was used as a measurement of the likelihood of randomly finding a node (specifically *A. muris* and *M. intestinale*) with the given BiBC and degree (or higher). A large value (i.e. a dark-colored space in the contour map) indicates that a node in that area typically occurs in random networks size-matched to the network generated from the data.

## Supporting information

All supplementary figures/tables

