## Supplementary material for "Host response to cholestyramine can be mediated by the gut microbiota": All supplementary figures/tables

**S1.** Experimental design for both experiments.

**S2.** Glucose levels 15, 30, 60, and 120 minutes after intraperitoneal glucose injection.

**S3.** PCoA plots comparing feature-level differences of ASVs between treatment groups.

**S4.** Order level stacked abundance plots.

**S5.** Comparisons of significantly changed bacterial families in cholestyramine-treated mice (Mann-Whitney U FDR < 0.05).

**S6.** The cholestyramine network follows a power-law degree distribution.

**Supplementary tables**

**ST1.** Mann-Whitney comparison results for each of the measured metabolic parameters. One-tailed p-values are reported for phenotypes and two-tailed p-values are reported for genes.

**ST2.** Family level comparisons between the WD+Ch and WD groups (two-tailed Mann-Whitney U test). Benjamini-Hochberg FDR correction is reported as well.

**S1**

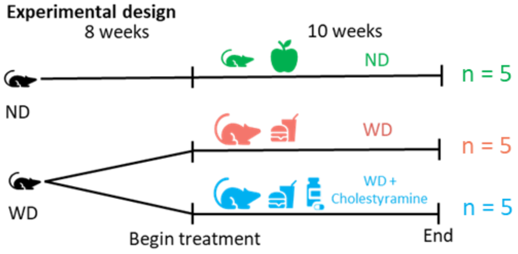


**S2**

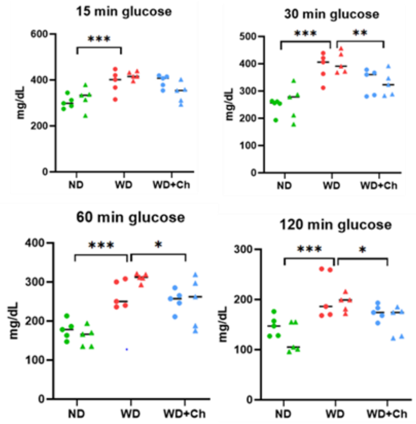


**S3**

## WD v ND

Experiment 1

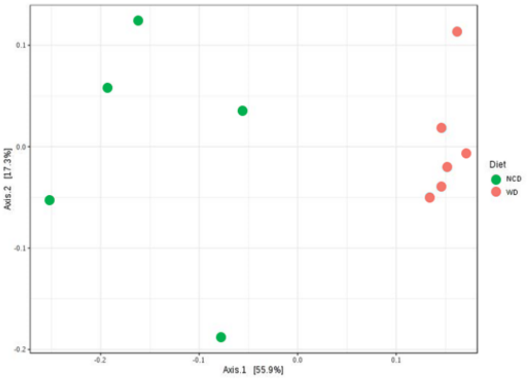


Experiment 2

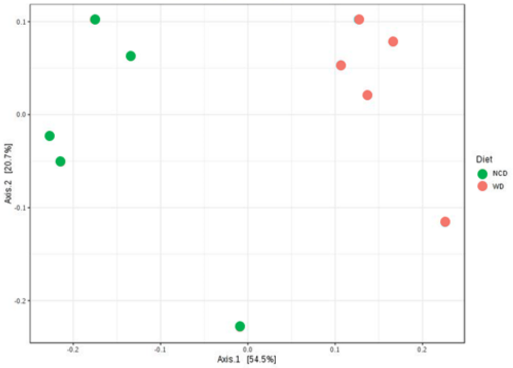


## WD+Ch v WD

Experiment 1

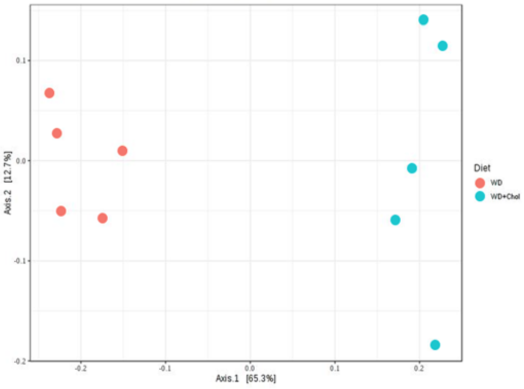


Experiment 2

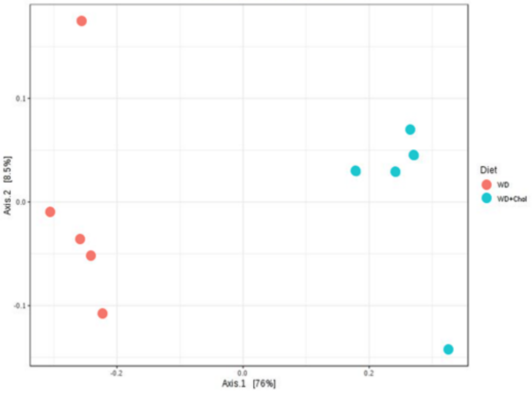

**S4**

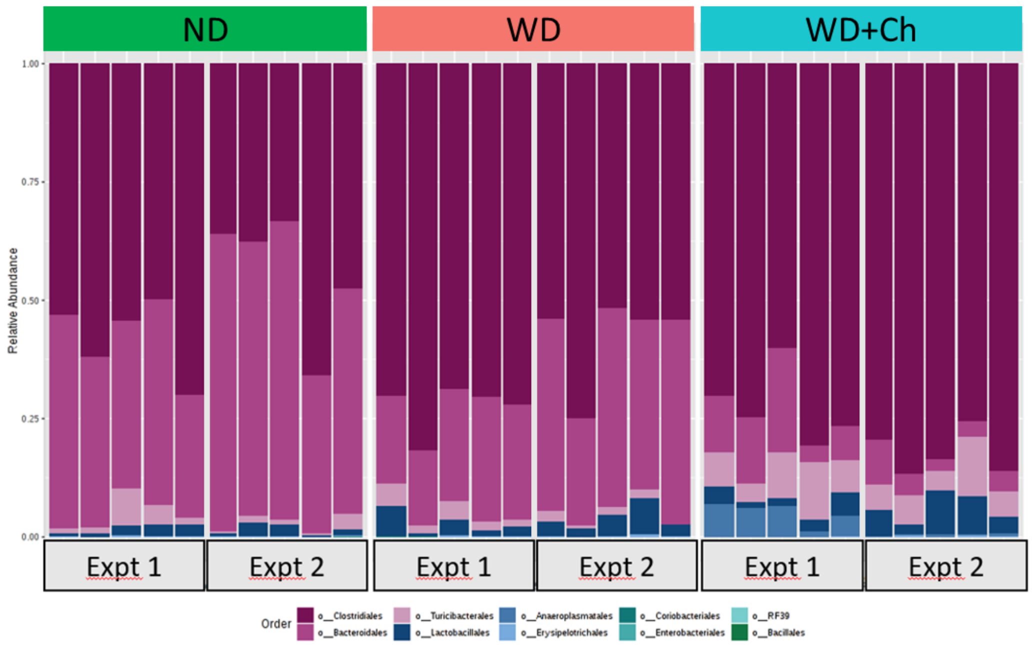
ND
WD
WD+Ch
Relative Abundance
1.00
0.75
0.50
0.25
0.00
Expt 1
Expt 2
Expt 1
Expt 2
Expt 1
Expt 2
Order
o__Clostridiales
o__Bacteroidales
o__Turicibacterales
o__Lactobacillales
o__Anaeroplasmatales
o__Erysipelotrichales
o__Coriobacteriales
o__Enterobacteriales
o__RF39
o__Bacillales

**S5**

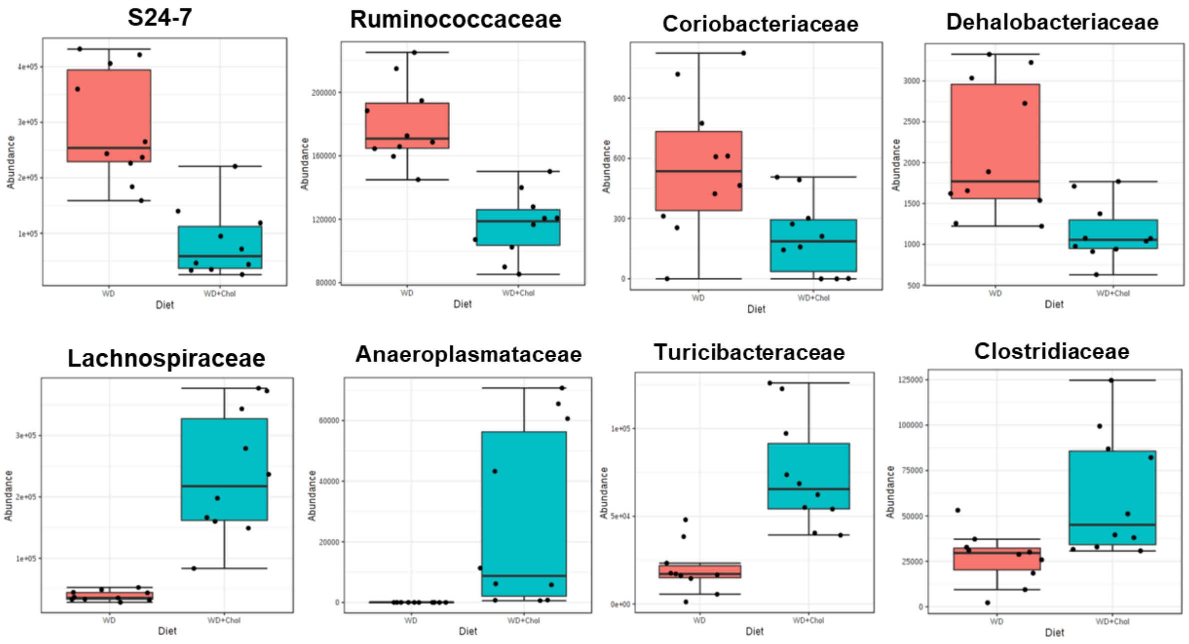
S24-7
Ruminococcaceae
Coriobacteriaceae
Dehalobacteriaceae
Lachnospiraceae
Anaeroplasmataceae
Turicibacteraceae
Clostridiaceae
Abundance
Diet
WD
WD+Chol

**S6**

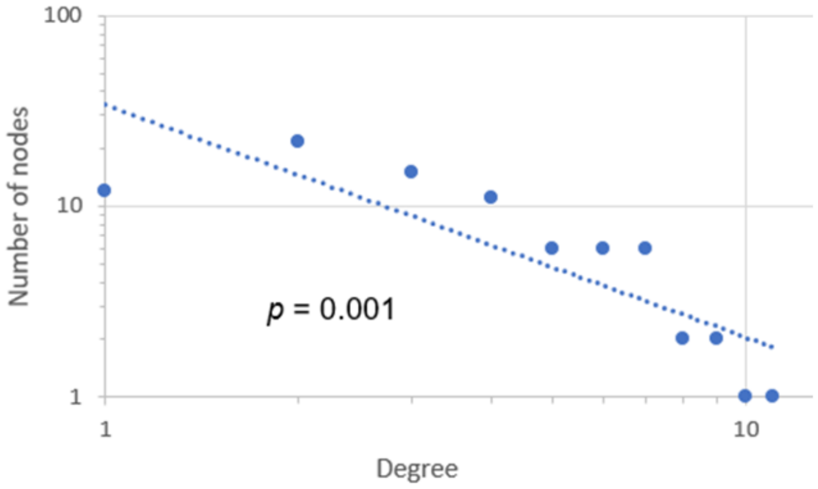


6) The cholestyramine network follows a power-law degree distribution.

**ST1**

| | Parameter | WD v ND | WD+Ch v WD |
|---|---|---|---|
| One-tailed p-value | Fed glucose | 0.0004 | 0.0014 |
| | Body weight | 0.0001 | 0.5000 |
| | Fasting glucose | 5.41E-06 | 0.0026 |
| | Glucose (15 min) | 0.0005 | 0.0205 |
| | Glucose (30 min) | 0.0001 | 0.0073 |
| | Glucose (60 min) | 0.0001 | 0.0319 |
| | Glucose (120 min) | 0.0002 | 0.0205 |
| | Fasting insulin | 0.0315 | 0.4399 |
| | GTT-AUC | 5.41E-06 | 0.0023 |
| | Serum cholesterol | 0.0003 | 0.1132 |
| | Epididymal fat | 0.0008 | 0.0086 |
| Two-tailed p-value | fgf15-ileum | 0.3256 | 1.08E-05 |
| | gcg-ileum | 0.8798 | 0.0068 |
| | shp-ileum | 0.3527 | 0.0001 |
| | fxr-liver | 0.0029 | 0.0892 |
| | shp-liver | 0.0433 | 0.0036 |
| | cyp7a1-liver | 0.1040 | 0.0002 |
| | tgr5-ileum | 0.2567 | 0.9118 |
| | tgr5-liver | 0.0288 | 0.5452 |
| | fxr-ileum | 0.6842 | 0.7394 |

## ST2

| Family | p-value | FDR | Change direction |
|---|---|---|---|
| Lachnospiraceae | 1.08E-05 | 0.000206 | Up |
| Ruminococcaceae | 2.17E-05 | 0.000206 | Down |
| S24_7 | 4.33E-05 | 0.000206 | Down |
| Turicibacteraceae | 4.33E-05 | 0.000206 | Up |
| Anaeroplasmataceae | 6.39E-05 | 0.000243 | Up |
| Clostridiaceae | 0.00288 | 0.007816 | Up |
| Dehalobacteriaceae | 0.00288 | 0.007816 | Down |
| Coriobacteriaceae | 0.018661 | 0.039395 | Down |
| Peptostreptococcaceae | 0.052426 | 0.090554 | NS |
| Paenibacillaceae | 0.077872 | 0.1233 | NS |
| Enterococcaceae | 0.12061 | 0.16694 | NS |
| Erysipelotrichaceae | 0.12301 | 0.16694 | NS |
| Mogibacteriaceae | 0.21756 | 0.27558 | NS |
| Staphylococcaceae | 0.26735 | 0.31748 | NS |
| Lactobacillaceae | 0.43587 | 0.48715 | NS |
| Enterobacteriaceae | 0.59531 | 0.62839 | NS |
| Streptococcaceae | 0.9118 | 0.9118 | NS |
